# Heterogeneity and ‘memory’ in stem cell populations

**DOI:** 10.1101/2020.09.22.307850

**Authors:** Patrick S. Stumpf, Fumio Arai, Ben D. MacArthur

## Abstract

Modern single cell experiments have revealed unexpected heterogeneity in apparently functionally ‘pure’ cell populations. However, we are still lacking a conceptual framework to understand this heterogeneity. Here, we propose that cellular memories – the ability of individual cells to record their developmental past and adapt their response to their environment accordingly – are an essential ingredient in any such theory. We illustrate this idea by considering a simple age-structured model of stem cell proliferation. Using this model we argue that heterogeneity is central to stem cell population function, and memories naturally explain why stem cell numbers increase through life, yet regenerative potency simultaneously declines.

## Introduction

Modern experimental methods allow us to profile the molecular state of hundreds of thousands of individual cells in a single experiment, and explore cellular identities in unprecedented detail (Shapiro et al. 2013, Stuart & Satija 2019). Accordingly, a number of efforts to catalogue the cellular diversity in multicellular organisms are now well under way, including the human cell atlas (Regev et al. 2017), the mouse cell atlas (Han et al. 2018), and maps of various model organisms including *D. rerio* (Wagner et al. 2018), *D. melanogaster* (Karaiskos et al. 2017), *C. elegans* (Cao et al. 2017), the planarian *S. mediterranea* (Plass et al. 2018, Fincher et al. 2018) and the cnidarian *N. vectensis* (Sebé-Pedrós et al. 2018). These efforts are important not only to build our understanding of healthy biology, but also to determine how different cellular identities, and the balance of cell types present in tissues and organs, change with disease and age (Regev et al. 2017).

Central to these efforts is the desire to properly understand the mapping from cellular molecular ‘states’ (i.e. patterns of gene and protein expression, or other regulatory molecules, within the cell) to cell ‘types’, ‘fates’ or functions. However, despite tremendous progress in this area, the regulatory principles that underpin this mapping are still unclear. Indeed, despite its centrality to our understanding of multicellular life, we still do not have a clear notion for what we mean by a cell ‘type’ (Clevers et al. 2017).

It is worth noting that until the pioneering work of Gurdon and co-workers in the 1970s it was not even apparent if each cell in the adult organism possessed the same genome, or only the portion of the genome it needed to perform its particular function (Gurdon et al. 1975). We now know that most adult cells possess essentially the same complement of genetic information (except certain lymphocytes, some neurons and mature red blood cells) (Allis et al. 2015), yet vastly different patterns of gene expression are observed between cells. This is a remarkable fact – and one that allows for the possibility that individual cells can be reprogrammed in a myriad of ways, thus paving the way for personalized regenerative medicine (Rackham et al. 2016) – the consequences of which we are still exploring.

Typically, cell types are defined informally based on experimentally observable features such as cell morphology, gene and protein expression patterns and the ability to perform specialised functions. Yet, both cellular form and function are complex issues that depend on the cell’s environmental context and it’s developmental history (Clevers et al. 2017). Furthermore, any practical definition of form, function or ‘type’ is contingent on what is experimentally observable, and some aspects of cellular character, such as developmental history, are hard to observe and harder still to quantify.

To deconvolute these issues, developmental biologists have traditionally mapped adult cell identities to cellular genealogies, since much of the cellular diversity observed in the adult organism is established during development (Davidson & Levin 2005). For example, distinct cell types are commonly associated with their germ layer of origin (Remak 1855). In eutelic organisms such as *C. elegans* (which contain a fixed number of cells at adulthood) we can go further, and whole organism lineage trees have long been used to trace the developmental histories of diverse adult cell types (Sulston et al. 1983).

Similar lineage trees have also been inferred in complex organisms such as zebra fish and mouse, using cell lineage tracing by imaging (Keller et al. 2008, McDole et al. 2018) and, increasingly, CRISPR/CAS9 induced genetic scar sequencing (Alemany et al. 2018, McKenna et al. 2016, Kalhor et al. 2018), amongst other methods (Kretzschmar & Watt 2012, Woodworth et al. 2017, Kester & van Oudenaarden 2018). These lineage maps help show how distinct cell types are produced during development in a regular and reproducible way. Lineage tracing studies using viral barcoding or genetic reporters have also helped elucidate the regenerative potential of individual stem cells and hence the mechanisms by which the numbers of cells under continuous turnover in the adult organism are maintained (Lemischka et al. 1986, Lu et al. 2019, Nguyen et al. 2014, Snippert et al. 2010).

From these studies it is becoming increasingly clear that many functional cell types are highly heterogeneous in the sense that cells in a variety of different molecular states (i.e. expressing different patterns of genes and proteins) may all perform the same function. Mathematically the mapping from cell states to cell types is many-to-one. This has an important consequence because it suggests that each cell is an individual and while they may perform ostensibly similar tasks, variability within cellular identities is common and, in some cases, may even be fundamental.

The role of mitotic memory in regulating the function of stem cell populations is a notable example. It has been observed that divisions of sister cells from hematopoietic stem cell (HSC) divisions soon fall out of sync with each other in vitro (Suda et al. 1984*b*,*a*), and individual HSCs can differ widely both in their cell cycle times and the rate at which they enter the cell cycle (Roch et al. 2017, Cheshier et al. 1999, Schroeder 2013). In some cases this variability may derive from heterogeneity in the population with respect to cell cycle rates (Fleming et al. 1993). However, this is not always the case. Broad variability in cell cycle times are expected to emerge naturally, even within ‘pure’ populations, due either to inherent stochasticity or complex determinism in the molecular mechanisms that drive cell divisions (Yates et al. 2017, Shields 1977, Sandler et al. 2015). During native hematopoiesis in vivo it is not, therefore, expected that HSCs will synchronize their cell cycles throughout life.

This lack of synchronization is important because if HSC divisions are temporally variable then different cells in the stem cell pool will accumulate a different number of divisions as the organism (e.g. mouse or human) ages, generating a population that is inherently heterogeneous with respect to mitotic history (see **Fig. 1**). In principle this mitotic heterogeneity need not be of functional importance. However, if the accumulation of prior cell divisions confers distinct biases to individual stem cells then mitotic history may be an important regulator of HSC function.

**Figure 1.**
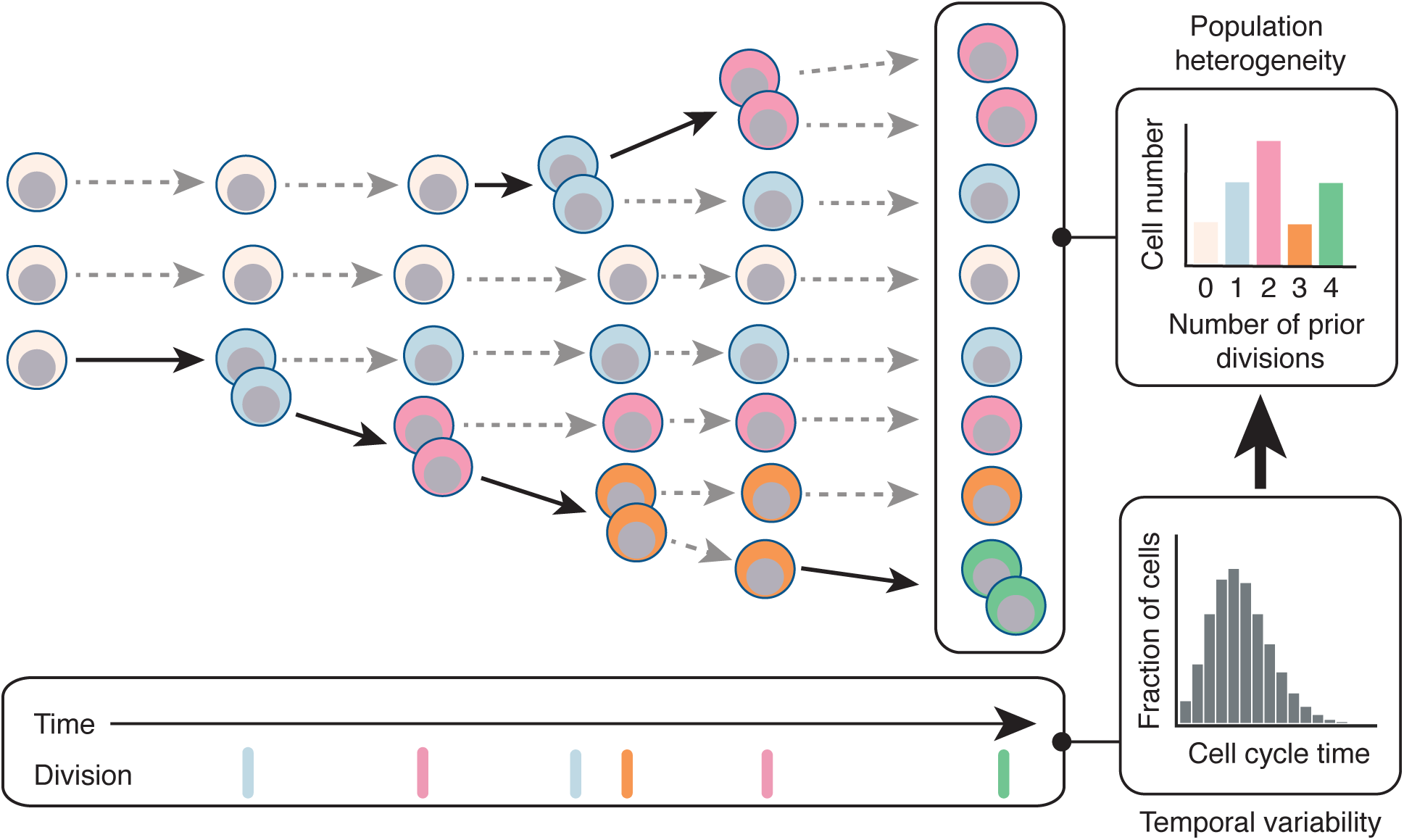
Cell proliferation produces heterogeneity. Starting from a homogeneous population of stem cells, variations in cell cycle rates naturally generate heterogeneity within the proliferating population in which individual cells are distinguished by their mitotic history. If mitotic history affects cell function then variations in cell cycle rates can produce a functionally heterogeneous population. Bold arrows indicate cell division events, dotted arrows represent perpetuation of cells in their current state. Cells are colored by number of prior divisions.

There is emerging evidence that this is the case. A number of recent reports have found that repopulation activity is concentrated in the most quiescent HSC subpopulations (Wilson et al. 2008, Foudi et al. 2009, Takizawa et al. 2011, Qiu et al. 2014) and, furthermore, the accumulation of cell divisions is directly associated with loss of stem cell potency both during native hematopoiesis and under conditions of stress (Bernitz et al. 2016, Säwén et al. 2016, Qiu et al. 2014, Wilson et al. 2008, Nygren & Bryder 2008, Foudi et al. 2009, Arai et al. 2019). Although the precise reasons for this association are unclear, it is likely that HSCs are sensitive to the general wear and tear that occurs during cell division (Walter et al. 2015). Cells naturally acquire genetic alterations on cell division, and proliferation can thereby give rise to complex mosaics (Ju et al. 2017) that can significantly affect HSC population function (Haas et al. 2018). General mechanisms, such as telomere shortening, may also define a ‘mitotic clock’ that places a fundamental limit on HSC proliferative capacity in vivo (Harley et al. 1990, Notaro et al. 1997, Hills et al. 2009, Allsopp, Morin, DePinho, Harley & Weissman 2003, Allsopp, Morin, Horner, DePinho, Harley & Weissman 2003, Werner et al. 2015). Remarkably, there is evidence that – however it is recorded – such a mitotic clock is encoded reliably enough that individual HSCs are able to ‘count’ cell divisions and thus have a ‘memory’ of their mitotic history (Bernitz et al. 2016).

Collectively these results indicate that mitotic memories play an important part in regulating stem cell population function. However, the precise role of mitotic memories is unclear. To explore putative roles for memories further, here we will consider a simple model of stem cell proliferation as an age-structured process. To make ideas concrete, We will frame our model in the context of hematopoiesis, but note that comparable age-structured processes may be play a central part in regulating stem cell dynamics in other contexts.

### An age-structured model of HSC proliferation

Age-structured processes arise whenever ageing affects the behaviour of individuals within a population, and age-structured models have accordingly been used to model a wide-range of biological and epidemiological processes (Li & Brauer 2008, Auger et al. 2008). At the beginning we should stress that our model is not intended as a detailed molecular description of stem cell proliferation, but rather a simple conceptual framework in which the implications of the connections between temporal cell cycle variations and heterogeneity can be explored.

To start we consider a population of HSCs proliferating under normal conditions in vivo and assume that stem cell divisions can be of three types: (1) symmetric S-S divisions, which produce two daughter stem cells; (2) asymmetric S-D divisions which produce one daughter stem cell and one differentiated daughter cell; and (3) symmetric D-D divisions, which produce two differentiated daughter cells.

To encode mitotic memory we allow the propensities for S-S, S-D and D-D divisions to depend on the number *n* of prior divisions that the stem cell has undergone. For simplicity, we will assume that all adult HSCs arise from a common population of founder cells during development – for example, hemogenic endothelial precursor cells (Bertrand et al. 2010) – and begin counting divisions at the onset of adulthood. To allow for niche regulation of stem cell divisions, we additionally assume that the local micro-environment of the cell is described by a single variable *m* that also affects cell divisions. Without loss of generality we can assume that 0 ≤ *m* ≤ 1 such that *m* = 1 implies a ‘strong’ niche that maximally stimulates self-renewal and *m* = 0 implies a ‘weak’ niche that does not support self-renewal. Thus, the variable *m* provides a simple way to take account of the numerous niche factors that collectively regulate stem cell divisions.

To account for evolving division patterns, let *p*_11_(*n*|*m*), *p*_10_(*n*|*m*) and *p*_00_(*n*|*m*) be the conditional probabilities that a stem cell that has divided *n* times previously will undergo S-S, S-D or D-D division respectively in the niche *m*. Thus, division probabilities are governed by a bivariate Bernoulli distribution, which may be written in general form as

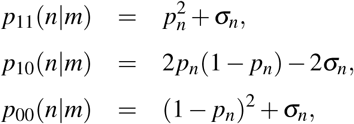

where *p*_*n*_ = *p*_*n*_(*m*) is the probability that each daughter cell will adopt a stem cell identity in niche *m* and *σ*_*n*_ = *σ*_*n*_(*m*) is the covariance between the daughter identities in niche *m*. If *σ*_*n*_ = 0 then daughter cell identities are adopted independently of each other; while if *σ*_*n*_ *≠* 0 then daughter cells are produced in regulated pairs.

Under this model stem cell numbers are described by the following set of ordinary differential equations

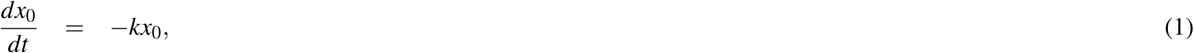

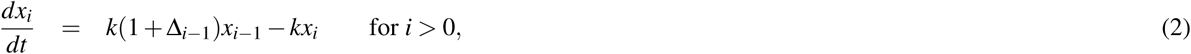

where *x*_*i*_ = *x*_*i*_(*t, m*) is the expected number of stem cells that have divided *i* = 1, 2 *…* times previously in niche *m*, Δ_*n*_ = Δ_*n*_(*m*) = *p*_11_(*n*|*m*) *− p*_00_(*n*|*m*) and ln(2)*/k* is the expected cell cycle time.

From these equations it is clear that the ability of a stem cell that has divided *n* times previously to self-renew in the niche *m* is quantified by the factor Δ_*n*_(*m*). In particular if Δ_*n*_(*m*) *<* 0 then D-D divisions predominate over S-S divisions and the niche is not strong enough to stimulate net self-renewal in the sub-population of cells that have divided *n* times previously (i.e. each cell division will produce less than one stem cell daughter on average). Conversely, if Δ_*n*_(*m*) *>* 0 then S-S divisions predominate over D-D divisions and the niche is able to net support self-renewal.

The total expected stem cell number *X* (*t*) = ∑_*i*_ *x*_*i*_(*t*) is governed by

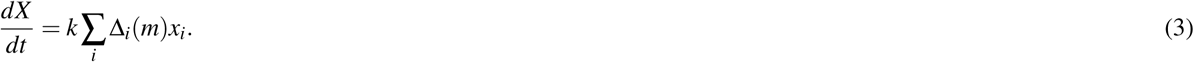

Thus, if Δ_*i*_(*m*) = 1 for all *i* then all stem cells in the pool undergo S-S divisions and the pool grows exponentially with time; if Δ_*i*_(*m*) = 0 for all *i* then *dX/dt* = 0 and the stem cell pool maintains homeostasis; and if Δ_*i*_(*m*) = *−*1 for all *i* then all stem cells in the pool undergo D-D divisions and the pool declines exponentially with time. These three special cases are not representative however, and in general the stem cell pool may expand or decline depending on the magnitude of

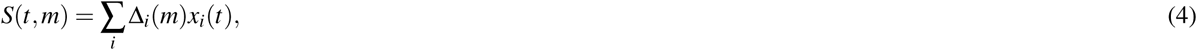

which represents the self-renewal ability of the population as a whole in niche *m*, taking into account its age-structure and the effect of mitotic history on each individual cell’s self-renewal ability. In particular, if *S >* 0 then the stem cell pool will be expanding, while if *S <* 0 then the pool will be in decline.

Notably, both stochastic and instructive dynamics are allowed in this framework, depending on the way that division history *n* and niche instruction *m* interact to co-regulate *p* and *σ*. For example, if *σ* = *p*(1 *− p*) *−* 1/2 then the niche unambiguously instructs all divisions to be asymmetric (of type S-D); if *σ* = *p*^2^ *−* 1/2 then the niche unambiguously instructs all divisions to be of type S-S; and if *σ* = (1 *− p*)^2^ *−* 1/2 then the niche unambiguously instructs all divisions to be of type D-D.

To make this idea more apparent, an explicit model for the interplay between division history and niche instruction is needed. From a mathematical perspective, functional forms for *p*_*n*_(*m*) and *σ*_*n*_(*m*) are required to close the model. In principle there many ways that the interplay with division history and niche instruction could affect cell divisions. Very little is currently known about this interplay except that division history diminishes self-renewal ability (Bernitz et al. 2016, Arai et al. 2019, Säwén et al. 2016). Since a simple model suffices to illustrate our point, we will therefore assume that daughter cells are produced independently of each other (i.e. *σ*_*n*_(*m*) = 0 for all *n, m*) and the likelihood that daughter cells adopt a stem cell identity declines with division history (i.e *p*_*n*_(*m*) is monotonic decreasing in *n*). For illustrative purposes we will assume an exponential loss of niche sensitivity with mitotic history and set *p*_*n*_(*m*) = *m*^*n*^. A schematic of this model is given in **Fig. 4A**.

### Model implications

Although clearly a simplification, this model illustrates how the complex interplay between stochastic and instructive mechanisms can arise in an age-structured proliferating stem cell population. It has a number of implications.

Firstly, within a defined niche (i.e. for 0 *< m <* 1 fixed), stem cell self-renewal ability will decline with number of divisions (see **Fig. 4B**). Juvenile stem cells will have a strong preference for symmetric S-S divisions that expand the stem cell pool. However, as individual stem cells accumulate cell divisions there will be a shift away from the S-S divisions that are characteristic of young cells and toward asymmetric S-D divisions that maintain but do not expand the stem cell pool. Finally, into old age it becomes increasingly difficult for the niche to maintain self-renewing divisions, and a bias toward D-D divisions begins to emerge (see **Fig. 4B-C**).

Secondly, the degree of stochasticity of stem cell divisions also changes with mitotic history. Because juvenile stem cells have a strong preference for S-S divisions, yet the balance between divisions shifts through mid-life and toward D-D divisions in later life, divisions of young and old stem cells are primarily instructive (i.e. unambiguously or strongly determined by the niche), while those through the middle of life are highly stochastic. The Shannon entropy is a simple information-theoretic way to quantify how stochastic an event is (Cover & Thomas 2012), and so can be used to quantify this shift (see **Fig. 4D)**. In this case, the division entropy *H*_*n*_(*m*) in niche *m* for a stem cell that has divided *n* times previously is

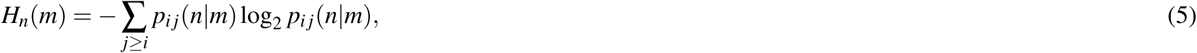

for *i, j* = 0, 1.

Thirdly, while stem cell division propensities and self-renewal abilities naturally change with mitotic history, these age-related changes can be continuously re-set by the niche. Particularly, aged cells can be ‘reactivated’ by transient guidance from a more strongly instructive niche. For instance, if *m* = 1 (i.e. cells are placed in a niche that is unambiguously instructive) then *p*_11_ = 1 regardless of *n*. Thus, all cells – even those that have accumulated a large number of divisions under native conditions and so have diminished self-renewal capacities – can be stimulated to exclusively undergo S-S divisions which rapidly expand the stem cell pool if needed. Such conditions may occur on conditions of stress, for example on bone marrow transplantation or after injury (Mendelson & Frenette 2014). Indeed, it has been observed that HSCs that have divided numerous times in vivo can be rapidly activated in this way (Wilson et al. 2008). Thus, this model accounts age-structured proliferation under conditions of homeostasis, yet allows for emergency hematopoiesis when it is needed (see **Fig. 4E**).

**Figure 2.**
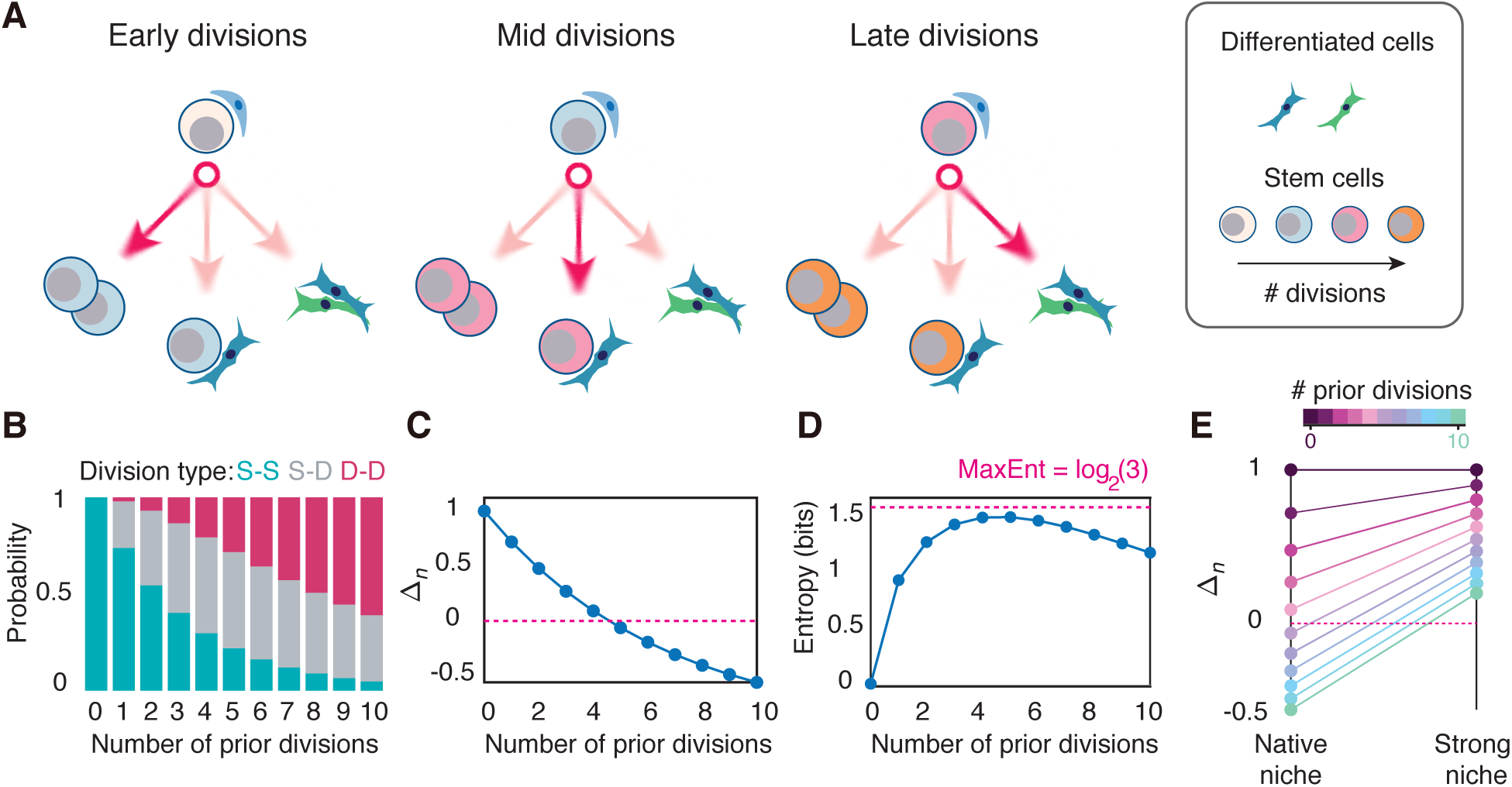
An age-structured model of HSC proliferation. (**A**) Schematic of model. Initially HSCs have a strong bias toward S-S divisions that expand the stem cell pool. As cell divisions accumulate there is a shift first toward asymmetric S-D divisions that maintain the stem cell pool and then toward D-D divisions that deplete the stem cell pool. (**B**) Division probabilities by mitotic history using *m* = 0.86, estimated from HSC proliferation data (see **Fig. 5**). (**C**) Within a given niche (i.e. for *m* fixed) the self-renewal potential of individual HSCs (Δ_*n*_ = *p*_11_(*n*) *− p*_00_(*n*)) declines with mitotic history. For *m* = 0.86 individual cells lose net self-renewal ability once they have accumulated four divisions (i.e. Δ_*n*_ *<* 0 for all *n* ≥ 5). (**D**) The stochasticity of divisions, as assessed by the Shannon entropy (Cover & Thomas 2012), is low for cells that have divided 0-1 times, reaches a maximum for cells that have divided 4-5 times, and begins to decline as cells accumulate numerous prior divisions. The maximum possible entropy, which occurs when all three division types are equally likely, is shown in pink. (**E**) Under normal conditions (*m* = 0.86) the self-renewal ability of individual HSCs is lost after 4 divisions. However, if cells are moved to a strong niche, for example under conditions that promote emergency hematopoiesis, then individual cells can be rejuvenated in their self-renewal ability. Here *m* = 0.95 is taken as a representative ‘strong’ niche.

### Implications for ageing

To investigate the relevance of this simple model we fitted Eq. 3 to long-term experimentally determined long-term repopulating stem cell numbers in the mouse (see Methods for experimental details). In general the model has two free parameters: *k*, the rate of stem cell proliferation and *m*, the strength of the in vivo HSC niche. It is well-known that the most potent HSCs divide only very rarely in vivo and estimates suggest that mouse HSCs divide only once every 145 days or approximately 5 times in a mouse lifetime (Wilson et al. 2008). Since this parameter is well-established, and to produce a parsimonious model, we will fix the cell cycle time at 145 days, and leave only the niche strength *m* as a free parameter. Although the resulting mathematical model Eq. 3 has only one free parameter, it provides a remarkably good fit to the data (see **Fig. 5A**), suggesting that despite its simplicity it may capture the essential characteristics of stem cell ageing.

In the context of ageing the dynamics described by Eqs. 1-2 naturally give rise to an evolving age-structured population in which each stem cell in the pool exhibits a different propensity for self-renewal depending on its particular mitotic history (see **Fig. 4E** and **Fig. 5B-C**). Importantly, this means that the potency of the population as a whole depends on the collective dynamics of an inherently heterogeneous mix of cells with different innate regenerative abilities (see **Fig. 4D**).

**Figure 3.**
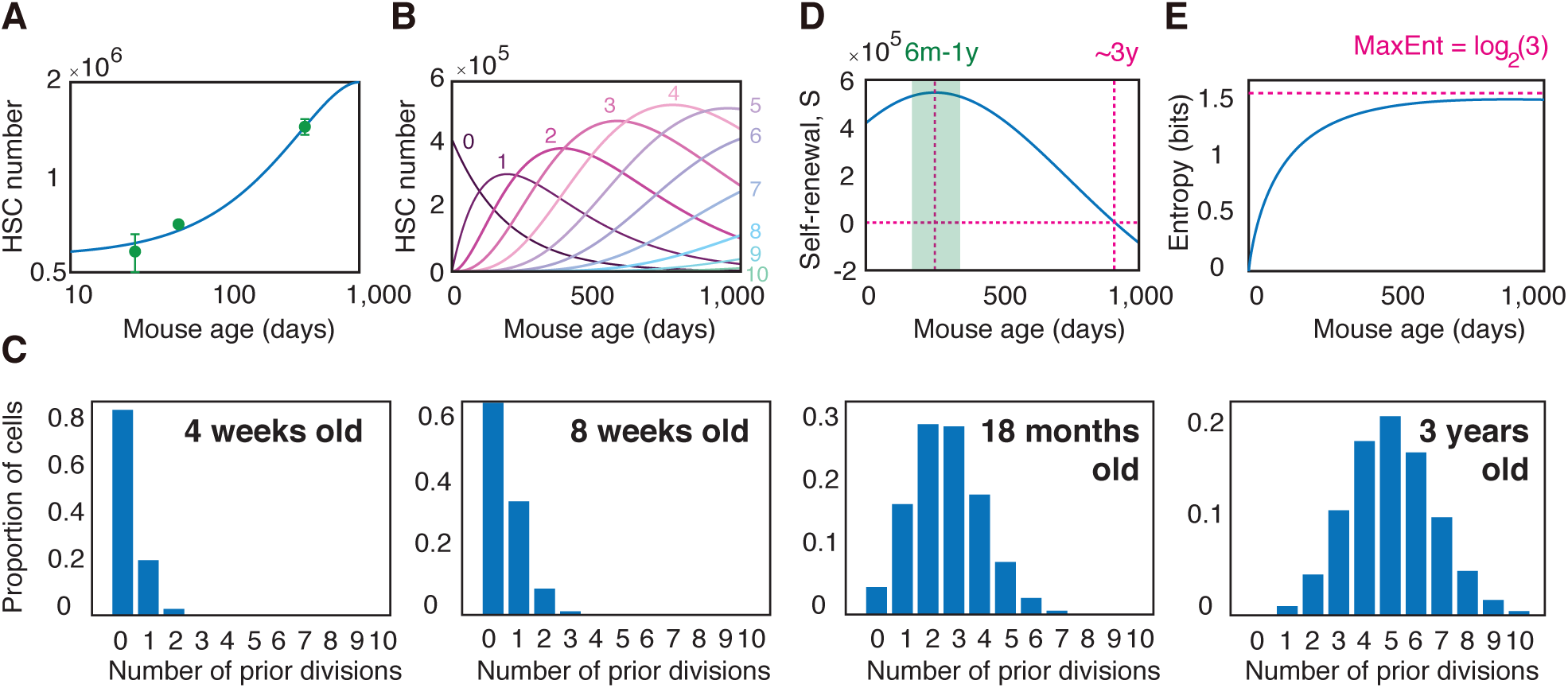
Changes in stem cell dynamics with age. (**A**) Fit of model (Eq. 3) to HSC numbers with age in the mouse. The model has only one free parameter *m*, which is estimated from data to be 0.86. A mean HSC cycle time of 145 days is assumed in accordance with experimental estimate (Wilson et al. 2008). See STAR Methods for details on experimental evaluation of HSC numbers. Experimental data is expressed as mean *±* standard deviation from three biological replicates. (**B**) Estimated HSC numbers over mouse lifetime as a function of number of prior divisions (0-10 shown). (**C**) Estimated proportion of cells in the HSC pool as a function of mitotic history. HSCs in young mice have typically not divided often, and the HSC pool is homogeneous with respect to mitotic history. As ageing occurs, individual HSCs accumulate different numbers of divisions and the HSC pool becomes heterogeneous with respect to mitotic history. (**D**) The self-renewal potential *S* of the HSC pool as a function of mouse age (i.e. the weighted average of Δ_*n*_). The HSC population reaches a maximum potency at age 6-12 months, and loses self-renewal ability at around 3 years. This estimate is in accordance with the approximate upper limit of natural mouse age. (**E**) The Shannon entropy of divisions as a function of mouse age (i.e. the weighted average of *H*_*n*_). Because the HSC population becomes increasingly heterogeneous as the mouse ages, the entropy of divisions also increases. In all panels *m* = 0.86 and a mean HSC cycle time of 145 days is taken.

Furthermore, in this age-structured setting, the distinction between stochastic and instructive divisions becomes opaque. Again the Shannon entropy can be used to assess division stochasticity. In particular, the weighted average

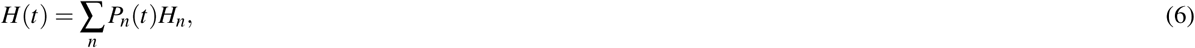

where *P*_*n*_(*t*) is the proportion of cells in the population that have divided *n* times previously, determines the stochasticity of divisions in the population as a whole, taking into account its mitotic heterogeneity. Analysis of this quantity indicates that generally more instructive dynamics will be observed in young individuals, with increasing stochasticity of divisions occurring with age (see **Fig. 5D**). However, at the individual cell level the view is more murky: depending on mitotic history some cells will respond unambiguously to niche instruction while others will divide more stochastically (see **Fig. 4E**). This issue will be compounded if niche conditions also vary (which is likely, see (Crane et al. 2017)). In this case, cells with the same number of prior divisions may behave more or less stochastically, depending on the particular niche within which they are located. Under these conditions it may be practically impossible, nor preferable, to attempt to dissect this complexity since it is an inherent part of how hematopoiesis is regulated. In this sense, stem cell proliferation is highly contextual, and both stochastic and instructive.

Finally, because the propensity for symmetric divisions that deplete the stem cell pool increases with age, in this model it is expected that the pool will exhaust on ageing if the stem cell population is too proliferative (note that in our simulations we used a fixed cell cycle time of 145 days, however in principle this parameter could also be varied). However, this does not appear to happen. Rather, it has been widely observed that the immunophenotypic stem cell pool increases with age, yet this increase is associated with loss of regenerative potential and increase in linage bias (Pang et al. 2011, Sudo et al. 2000, Rossi et al. 2005, Morrison et al. 1996, Geiger et al. 2013). Thus, maintenance of a viable stem cell pool with age requires that stem cells must either: (1) divide rarely, or (2) enter into a state of protective dormancy once their self-renewal ability has been sufficiently compromised (e.g. once Δ_*n*_ *<* 0). Thus, the notion that there must be a slowly cycling or dormant stem cell population arises naturally as a logical consequence of proliferation within an age-structured population. Both these options are possibilities. As already noted, it is well-known that the most potent HSCs divide only very rarely in vivo (Wilson et al. 2008). Similarly, it has been proposed that the most potent long-term repopulating HSCs divide only 4 times before entering a state of permanent protective dormancy (Bernitz et al. 2016, Wilson et al. 2008). It is currently unclear if this is hard limit or simply reflects the fact that most stem cells do not have time to divide more than four times within a single lifetime. Our model framework is consistent with either possibility.

## Conclusions

We have argued that mitotic memories shape the ability of individual stem cells to respond to niche direction and therefore have an important role in regulating stem cell fates. More generally, cellular memories – broadly defined as changes in the molecular status of a cell in response to a stimulus, that modify the ability of the cell to respond to future stimuli – may have a central role in fate regulation in numerous other contexts (Yu et al. 2016). Epigenetic mechanisms such as DNA methylation and histone modifications, are well-known to allow historical events to be encoded in stable, heritable modifications to the DNA and chromatin (Holliday & Pugh 1975, Razin & Riggs 1980, Bird 2002, Schübeler 2015, Allis & Jenuwein 2016, Gaydos et al. 2014). Remarkably, this memory storage can be sufficiently robust that it allows developmental programs to be reinstated in cells from epigenetic memory, thereby providing a ‘fossil record’ of development (Jadhav et al. 2019). Such epigenetic records are likely to play an important role in cell fate maintenance, regulation of differentiation, healthy aging, and malignant transformation in many situations (Yu et al. 2016, Hu & Shilatifard 2016, Haas et al. 2018, Challen et al. 2014, Trowbridge et al. 2009). Furthermore, it is also well-known that epigenetic alterations can confer substantial molecular and functional heterogeneity to otherwise ‘pure’ cell populations (Angermueller et al. 2016, Cheow et al. 2016). In light of this evidence, it is likely that individual cells commonly carry complex personal sets of epigenetic alterations that collectively confer subtle biases to their behaviour that are hard to quantify. Thus, each cell, like each person, is an individual, and while they may perform ostensibly similar tasks, variability within cellular identities is common and may indeed be fundamental. Consequently, the traditional strategy of organizing cells into discrete functional ‘types’ while undoubtedly practically useful, may ultimately be limited in its explanatory power since it does not fully acknowledge the subtle fecundity of cell biology. The difficulty is how to interpret such inherent cell-cell variability and to determine when it is functionally important and when it is not (Altschuler & Wu 2010).

Because they may be encoded in subtle, combinatorial ways, memories are hard to measure and dissecting their effect on cell fates is a significant technical challenge. Advances in experimental methods that track the mitotic history (or more generally the epigenetic history) of individual cells will help elucidate the mechanisms by which functional memories are formed and propagated. We anticipate that live-cell imaging, genetic barcoding and fluorescent label retention strategies (Kester & van Oudenaarden 2018, Schaniel & Moore 2009), in combination with the advanced mathematical and statistical models – including the latest advances in machine learning (Buggenthin et al. 2017, Arai et al. 2019) – needed to analyze the resulting data, will be particularly important in this regard.

## Methods

### Mice

Evi1-GFP knock-in mice were provided by Dr. Kurokawa (The University of Tokyo, Japan). The gene recombination experiment safety and animal experiment committees at Kyushu University approved this study, and all experiments were carried out in accordance with the guidelines for animal and recombinant DNA experiments.

### Flow cytometry and antibodies

The following monoclonal antibodies (Abs) were used for flow cytometry and cell sorting: anti-c-Kit (2B8, BD Biosciences, 1:100), -Sca-1 (E13-161.7, BD Biosciences, 1:100), -CD41 antibody (MWReg30, eBioscience, 1:100), -CD48 (HM48-1, Biolegend, 1:100), -CD150 (TC15-12F12.2, Biolegend, 1:100), and -CD34 (RAM34, BD Biosciences, 1:20). The following antiboodies were used in combination for the depletion of lineage-positive cells: anti-CD4 (RM4-5, BD Biosciences, 1:100), -CD8 (53-6.72, BD Biosciences, 1:100), -B220 (RA3-6B2, BD Biosciences, 1:100), -TER-119 (BioLegend, 1:100), Gr-1 (RB6-8C5, BD Biosciences, 1:100), and -Mac-1 (M1/70, BioLegend, 1:100).

### Cell isolation

Long-term repopulating HSC (LT-HSCs) were identified as Evi1-GFP+ LSKCD41-CD48-CD150+CD34-cells in 4, 8, and 60 week-old mice. To isolate Evi1-GFP+ LT-HSCs, Evi1-GFP mouse bone marrow cells were purified using a two-step protocol. First, LT-HSCs were enriched by positive selection for c-Kit expression using anti-CD117 immunomagnetic beads (130-091-224, Miltenyi Biotec Inc. 1:5 dilution). Second, c-Kit+ enriched cells were stained with a set of fluorophore-conjugated antibodies (lineage mix, anti-Sca-1, anti-c-Kit, anti-CD41, anti-CD48, anti-CD150, and anti-CD34). Estimates of total LT-HSC numbers were then calculated based on the total number of bone marrow mononuclear cells, frequency of LT-HSCs, and frequency of Evi1-GFP+ cells.

### Model fitting

Fitting of LT-HSC numbers to Eq. 3 was performed by minimizing the residual sum of squares between the experimental data and the model. Optimization was performed using the Nelder-Mead simplex algorithm, implemented in MATLAB® version 2018b (The MathWorks Inc., Natick, MA).

## Notes

### Competing Interest Statement

The authors have declared no competing interest.

